# Enabling population assignment from cancer genomes with SNP2pop

**DOI:** 10.1101/368647

**Authors:** Qingyao Huang, Michael Baudis

## Abstract

For a variety of human malignancies, incidence, treatment efficacy and overall prognosis show considerable variation between different populations and ethnic groups. Disentangling the effects related to particular population backgrounds can help in both understanding cancer biology and in tailoring therapeutic interventions. Because self-reported or inferred patient data can be incomplete or misleading due to migration and genomic admixture, a data-driven ancestry estimation should be preferred. While algorithms to analyze ancestry structure from healthy individuals have been developed, an easy-to-use tool to assign population groups based on genotyping data from SNP profiles is still missing and benchmarking for the validity of population assignment strategy for aberrant cancer genomes was not tested.

We benchmarked the consistency and accuracy of cross-platform population assignment. We also demonstrated its high accuracy to process unaltered as well as cancer genomes. Despite widespread and extensive somatic mutations of cancer profiling data, population assignment consistency between germline and highly mutated samples from cancer patients reached of 97% and 92% for assignment into 5 and 26 populations respectively. Comparison of our benchmarked results with self-reported meta-data estimated a matching rate between 88% to 92%. Despite a relatively high matching rate, the ethnicity labels indicated in meta-data are vague compared to the standardized output from our tool.

We have developed a bioinformatics tool to assign the populations from genome profiling data and validated its performance in healthy as well as aberrant cancer genomes. It is ready-to-use for genotyping data from nine commercial SNP array platforms or sequencing data. This tool is effective to scrutinize the population structure in cancer genomes and provides better measure to integrate genotyping data from various platforms instead of self-reported information. It will facilitate research on interplay between ethnicity related genetic background and molecular patterns in cancer entities and disentangling possible hereditary contributions.

The docker image of the tool is provided in DockerHub as “baudisgroup/snp2pop”.

## Introduction

Cancer arises from the accumulation of genomic aberrations in dividing cells of virtually all types of proliferating tissues (somatic variations). The irregular cellular expansion and other hallmarks of cancer can result from a plethora of mechanisms affecting multiple cellular processes (1). Some oncogenetic pathways can be initiated by exogenous factors, e.g. tobacco smoke or ultraviolet radiation (2). However, exposure to carcinogenics varies in its effects for people from different genetic background, which suggests that somatic mutations can be influenced by inherited (“germline”) variations (3, 4).

Cancer studies have reported significant world-wide variation in incidence and prognosis between ethnicity groups (5–8). While such differences have been attributed to unequal social and economical circumstances which influence risk factors and therapeutic interventions, several studies have shown impact of population specific genomic variants with predisposing effects on malignant transformation and phenotypic behaviour(9–12). Due to the late onset of most cancers, even high-penetrance Mendelian-type variants may not be purged by natural selection and can accumulate in particular populations. Such variants may play key roles in cancer development (13). Notably, mutations on BRCA1/2 genes confer a high risk to develop breast and ovarian carcinomas. Three founder mutations in Ashkenazi Jewish population cause the BRCA1/2 mutation prevalence to be 10-fold higher than all sporadic mutations in the general population (14, 15). Mitochondrial aldehyde dehydrogenase (ALDH2) encodes an enzyme in alcohol metabolism. Its “oriental” variant with 36% prevalence in East Asians, ALDH2*504Lys, increases risk for alcohol-related liver, colorectal and esophageal cancer by alcohol consumption (16, 17).

Many other studies have reported prevalent genetic variants in specific population groups which may contribute to the “racial” disparities in occurrence and prognosis (18–20). Other than these monogenic determinants, polygenic variation models for breast cancer which estimate the combined effect of multiple loci to be highly discriminatory in risk assessment, suggest the benefits of exploring genome-wide risk profiles (21). The potential impact of understanding the germline background of cancer genomes has also been demonstrated in a study which identified disease-associated chromosomal regions from only seven individual samples by using genome-wide relatedness or linkage mapping (22). This type of studies can be conducted population-wise, with sufficient number of samples from the same population or ethnicity group.

With the increasing number of available genome profiles and the decreasing cost to genotype clinical samples, the stratification between patients’ genetic backgrounds has become feasible with the promise to guide therapeutic strategies and improve the clinical prognoses. Since several studies have demonstrated the relevance of considering an individual’s genomic origin for preventive screening (reviewed in Foulkes *et al.* 2015 (15)), information about the population background of cancer patients may be an additional factor for individual therapeutic decisions as well as for the stratification of clinical study cohorts. A meta-analysis addressing the interplay between genetic background, cancer development and therapeutic responses is desirable, not only for robust statistical associations in molecular target identification, but also for the rational design of studies incorporating informative biosamples.

For many data repositories, “population group” of a sample can be assumed based on a geographical location associated with the sample. Alternatively, A self-reported “race” category is commonly used in U.S. census data. A biosample’s geographic origin is often approximated using the location of the study’s research facility or the contact address of its main authors. However, while these data can be easily retrieved, they may not provide an accurate representation of patients’ origins for the purpose of population-specific ancestry mapping. Self-reported data is often inconsistent across studies, vague in category description (e.g. *“white”*, *“black”* v.s. *“Caucasian”*, *“African”*) and misleading when patients have incomplete awareness of the migration and admixture histories of their ancestors. Overall, when associating oncogenic molecular signatures with germline variations, information from the above sources lacks in relevant detail and consistency.

A better approach to population assessment would be the computational estimate of ancestry with population-specific genomic variants. This has been shown previously for germline profiles, achieving 90% accuracy to distinguish three populations, African American, Asian and Caucasian, by using as few as 100 population-diverging single nucleotide polymorphisms (SNPs) (23), and nowadays is a standard methodology with claimed better granularity behind a number of commercial “ancestry” services. We hypothesise that a similar strategy can be applied to cancer genome data, despite the additional cancer-related somatic mutations which leads to both information loss (e.g. large scale homozygous or allelic deletions) and added noise (e.g. somatic mutations masking germline variants). An example of a cancer genome containing copy number loss and copy-neutral loss of heterozygocity (CN-LOH) events and its paired normal sample is shown in Figure 1. It is a typical case of altered genome with copy number loss (blue arrow) with CN loss regions partly recovered by doubling of the second allele (CN-LOH, black arrow) at multiple genomic locations. In addition to a general test of feasibility of inferring population background from the noisy cancer genome data, we continue with benchmarking population mapping procedures for heterogeneous datasets from different genotyping platforms, with the aim of integrating cross-platform cancer genotyping data for metaanalysis.

**Fig. 1.**
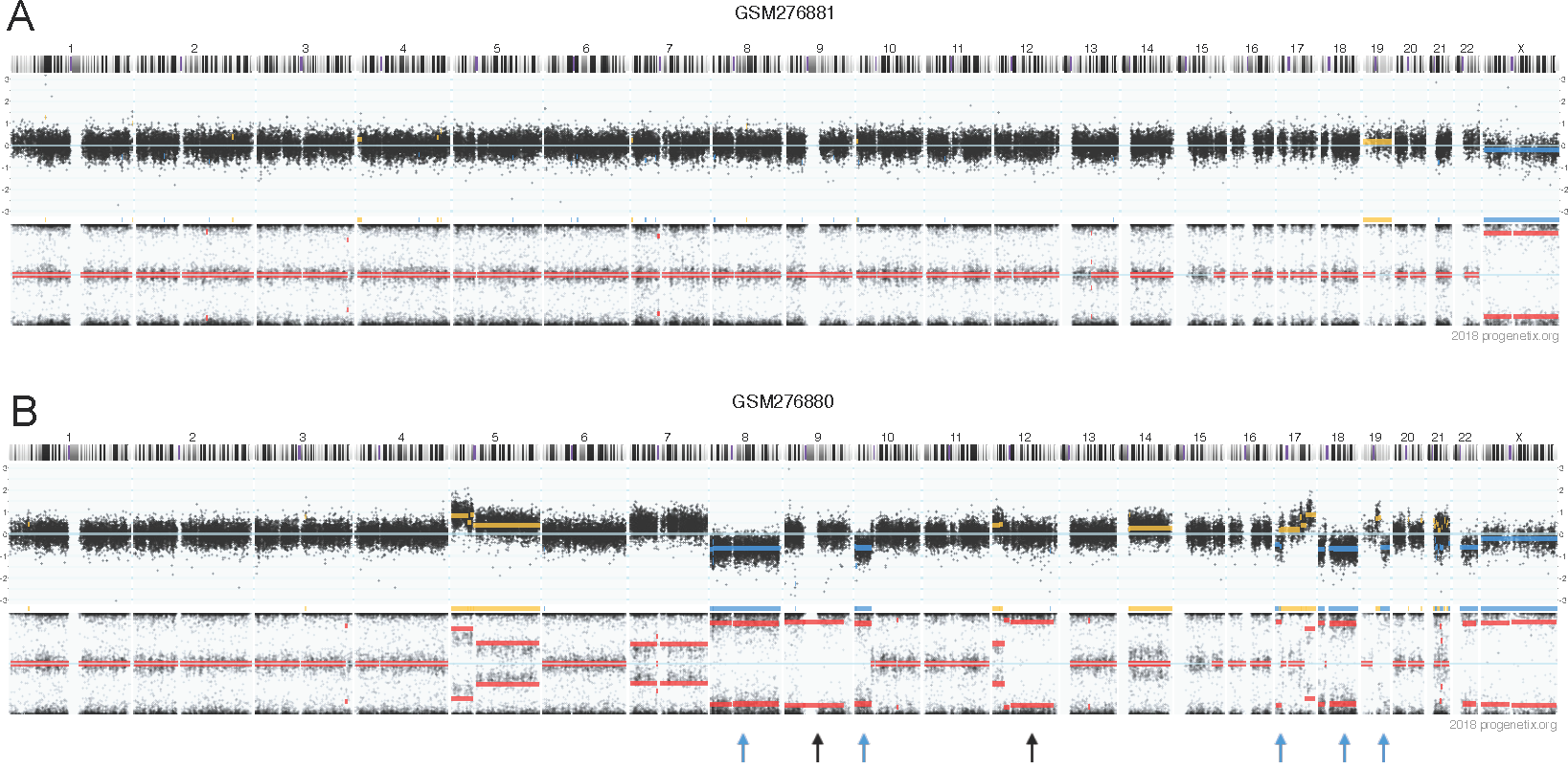
CNV tracks for a paired normal/cancer samples deposited in GEO with sample IDs GSM276881 and GSM276880. GSM276880 is a glioblastoma sample.GSM276881 is its peripheral blood control. The CNV tracks are consisted of two panels. Upper panel indicates the total copy number, represented by logR ratio (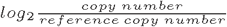 at any probe) Namely, logR = 0 indicates a position with normal two copies, logR > 0 indicates a copy number gain and logR < 0 indicates a copy number loss. Lower panel indicates the allele specific copy number, represented by B-allele frequency 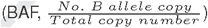. Any given SNP position can have a value between 0 and 1. A line at 0.5 indicates a heterozygous region. Compared to the unaltered genome (A), the cancerous counterpart (lower) has copy number loss in chr8, 10p, 18, 19qter, 22q and copy-neutral loss of heterozygocity in chr9,12q, accounting for 18.2% of genome.

## Methods

We retrieved the genomic reference data from the 1000 Genomes Project (24). SNPs of the selected array platforms were extracted from the sequencing data of these reference samples. In order to achieve between-study consistency for selection of informative SNPs, we used a model-based approach (25) where an admixture model is optimized with the reference set for each genotyping platform. The allele frequency and ancestry fraction parameters were projected to the incoming cancer dataset of the same platform. Applying a random forest classification, we assigned the population label to the highest voted group and produced a score for the difference between highest and second highest percentage votes. The overview of the pipeline is shown in Figure 2. The tool is accessible for direct use as a Docker image “https://hub.docker.com/r/baudisgroup/snp2pop” on DockerHub and its source codes can be found on Github in project “https://github.com/baudisgroup/snp2pop”.

**Fig. 2.**
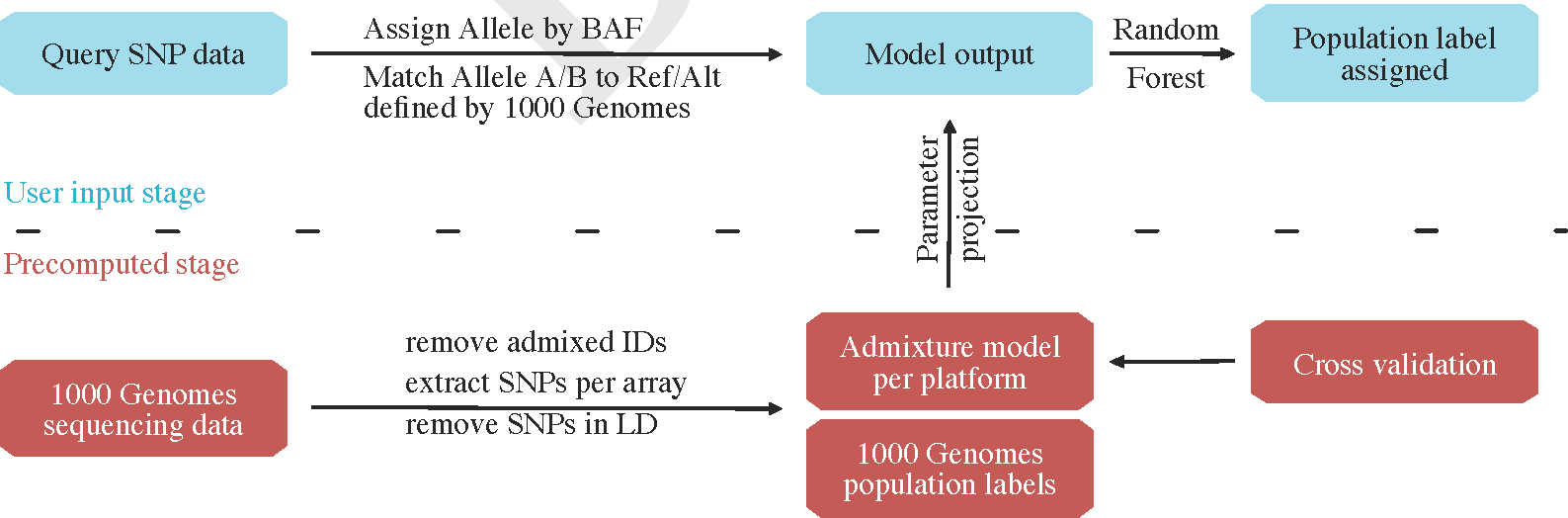
Pipeline of our tool to derive population assignment for individual cancer samples. The development workflow of the tool starts with retrieving the sequencing data from 1000 Genomes Project, removing IDs with admixture origin, extracting SNPs according to the genotyping array platforms and removing SNPs in linkage disequilibrium (LD). The generated PLINK files are stored as a intermediate reference file set. This file set is used further to train an admixture model with the model parameter matrix (allele frequency and ancestry fraction) stored for reference. Finally, a random forest classification is trained for the reference data based on their annotated population labels. These pre-computation steps are the basis of the tool, which saves time and preserves inter-study consistency. Now, when the new genotyping data comes in, the tool will assign the allele by B-allele frequency (BAF), check for the reference/alternative base for each SNP to match the reference data, assign the missing alleles (homozygous SNP from incoming data, thereby alternative base not available) based on reference SNP annotation. When the data is cleaned and compatible, the tool will use the pre-computed model parameters to analyze the admixture in the incoming dataset. Finally, with the model output, the population group of the incoming data is assigned. Users can choose to output the model parameters as well as the population groups.

### Data preparation

Reference sequencing data were provided by 1000 Genomes Project, a publicly available reference catalogue of human genotype variation (24). It sequenced 2504 individuals from 26 population groups out of 5 super population (continental) groups. With sequencing and subsequent multi-sample genotype imputation approach, the project covered human genetic variation in the studied population groups.

#### Reference data preparation

The SNP positions for each platform were acquired from Affymetrix annotation files. The allele information was extracted for all positions with vcftools (26). The 12 mislabeled or admixed individuals were removed from the reference dataset, leaving 2492 individuals. The SNP positions with duplicated rsIDs in annotation files were removed. The reference or alternative strands were swapped to match the SNP array annotation. Sites with minor allele frequency (MAF) of less than 5 percent were removed. SNPs were subsequently pruned for independence using PLINK 1.9 (27). Specifically, a sliding window of 50bp and a 5bp shift of window at each step of pruning, with the variance inflation factor (*VIF* = 1/(1 − *R*^2^), with *R*^2^ equal to the multiple correlation coefficient for a SNP regressed on all other SNPs) at 1.5. The result files were stored as PLINK output for each platform in ․bed, ․bim and ․fam formats, of which the ․bim files were used to extract SNP positions from target data.

#### Target data preparation

The SNP array data were processed with ACNE R package (28) to extract allele-specific copy numbers as B-allele frequencies (BAF). SNPs were labeled as homozygous A, heterozygous AB or homozygous B by the BAF value in ranges 0-0.15, 0.15-0.85 or 0.85-1, respectively, to allow both for noise and expected aneuploidy in the biosamples.

Data used for benchmarking are accessed through arrayMap database (29), using a collection of re-processed genotyping series from the Gene Expression Omnibus (GEO) repository (30), and the TCGA data repository (31) from https://cancergenome.nih.gov. The samples deposited in the former were submitted by individual research studies, whereas the latter coordinated cancer genome data generation and processing with centralized protocols.

### Admixture model

While many approaches use principle component analysis (PCA) to select informative SNPs for population assignment, deriving them prior to clustering methods, either by removing correlated SNPs (with Pearson’s r > 0.99) or by global fixation index (Fst > 0.45), results in varying sets of SNPs between datasets. Here, we used the allele information output from the reference panel to generate an admixture statistical model (25), which estimates the contribution of each SNP to the population category by alternately updating allele frequency and ancestry fraction parameters. Model was built based on two main parameters, the allele frequency of each SNP in the theoretical ancestors, and the fraction of theoretical ancestors in every sample. The number of theoretical ancestors(K) was defined by reference data with cross validation. It was chosen to be 9 as higher K did not further lower the cross-validation error. The ancestry fraction plot for reference individuals demonstrates a proper information extraction to distinguish the five continental categories. By projecting a correspondingly learned model derived from the reference dataset to a new sample with the corresponding platform, a robust and consistent output with 9 ancestry fractions was generated. The K fraction ratio of each individual in the reference data of 26 population groups for each platform is shown in Supplementary Figure S1.

### Random Forest label assignment

The nine ancestry fractions from the reference population were used to build a random forest model in each platform to predict the 26 population categories. The score was calculated as the difference in percentage votes between the best and the second best predicted labels. We performed repeated cross validation experiments (5-fold cross validation with 10 repetitions), and discovered twelve individuals assigned into a category different from the labels defined in their meta-data, which might occur due to a platform-specific detection of rare alleles that segregate in these samples. They were removed from reference list of the model. Finally, 2492 samples were used as training set in the classification model (removed IDs and information are found in Supplementary table S1). We compared random forest (RF) learning method with a classical multinomial linear regression model in terms of performance and computation time. The computation time is 3-5 fold slower than MLR (Supplementary Table S2) but the performance in terms of accuracy is moderately higher (Supplementary Figure S2). The classification strategy used in the tool is first getting votes for all 26 population groups, and generate a highest vote group as the prediction result. Then, users have options to further generate a prediction out of 5 superpopulation groups or 10 broader population groups as described in Section “Cross-platform benchmarking”, combining votes of groups belonging to the same super-population or broader population group, and output the group with highest vote as the prediction outcome, and a score which allows for closer scrutiny for potential admixtures.

## Results

We benchmarked our method with various normal and cancer samples from independent datasets to demonstrate the feasibility and reliability of this approach.

### Cross-platform benchmarking

We first used the original data from 1000 Genomes to validate the level of resolution needed for accurate population assignment from the pipeline. Taking the sequencing data of 2504 samples and extracting the SNPs from the nine genotyping platforms of interest gave rise to the dataset in this benchmarking experiment. The number of SNPs per platform ranged from 10 204 (Affymetrix Mapping 10K) to 934 946 SNPs (Affymetrix Genome Wide SNP 6). For all nine genotyping platforms (of seven levels of resolution), the model performed equally well in capturing the informative SNPs and predicting the population category. The assignment of 2504 samples from 1000 Genomes Project into 5 continental groups defined by the Project had low margins of error for all genotyping platforms (Figure 3). To benchmark the separation of 26 groups, we applied a repeated cross validation (CV) on both random forest and multinomial linear regression model for the assignment task. Here, the CV accuracy rate ranged between 75 – 85% depending on platform resolution. When combining pairs of frequently mixed groups into 10 categories, a CV accuracy rate rose to >99% (Supplementary Figure S3 and Supplementary Table S3).

**Fig. 3.**
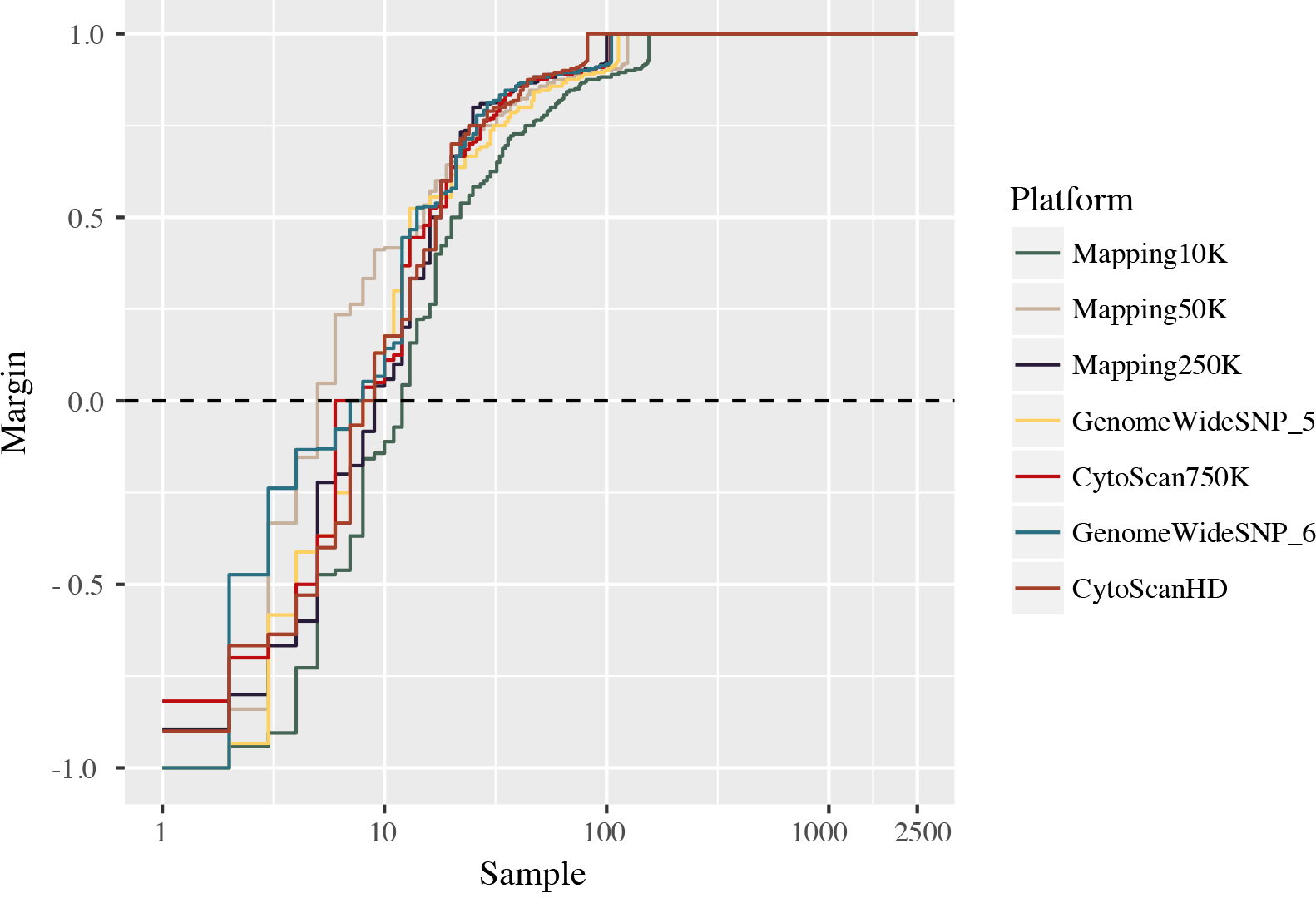
Estimating the model accuracy with reference samples with SNPs extracted from genotyping platforms of seven different resolution levels. Here we use margin as an estimator for the accuracy of the prediction model with random forest classifier output. It is defined as difference between the proportion of votes for the correct class minus maximum proportion of votes for the other classes. A positive value indicates correct prediction. For 2504 samples in the 1000 Genome Project, the number of incorrectly predicted samples ranged from 4 to 11, corresponding to an error rate of 0.16% to 0.4%. Note that this is cross validation error on reference dataset, rather than real data.

### Benchmarking normal genome profile assignment with HapMap data

To validate the accuracy of our tool to map population origins from non-cancer SNP datasets, we used 112 samples from the HapMap project (32) which is not included in the reference set, but acquired through GEO. The 112 samples came from three distinct population groups (at the level of 26 population groups, from three different super population groups): CEU (from EUR), CHB (from EAS) and YRI (from AFR). Except for five samples with “CEU” label predicted as “AMR”, the assignment of the rest of samples all matched their ethnicity information indicated in the metadata (Table 1).

**Table 1.** Comparison of HapMap metadata and predicted population group. Each column indicates HapMap population labels. Each row indicates the label predicted by the tool. CHB stands for Han Chinese in Beijing, China. YRI stands for Yoruba in Ibadan, Nigeria; AFR stands for African; AMR stands for Admixed American; EAS stands for East Asian; EUR stands for European.

|  | CEU | CHB | YRI |
| --- | --- | --- | --- |
| AFR | 0 | 0 | 45 |
| AMR | 5 | 0 | 0 |
| EAS | 0 | 6 | 0 |
| EUR | 56 | 0 | 0 |

### Paired cancer-normal comparison

One of the highlights in this tool lies in the determination of population origin from cancer genome profiles carrying various acquired genomic aberrations. Since the non-cancer samples could be correctly assigned according to HapMap categories, we then validated the cancer genome based assignments in samples where normal genome profiles of the same patients (e.g. from peripheral blood or non-cancer tissue samples) were available as reference. We performed the validation with two independent data sources — GEO and TCGA project.

#### GEO data

From the GEO repository, we retrieved all paired normal and cancer samples from 1145 individuals and compared the outcome of the population assignment. When including all 1145 individuals, 92.1% of the normal samples matched with paired tumor samples. With an increasing confidence score, the matching accuracy increased (Supplementary Table S4). The error rate dropped from 15.9% to 1.5 % comparing samples with score range “0.2-0.4” to those with “0-0.2”. After setting a threshold of normal samples with score > 0.2, 95.8% accuracy could be achieved for the remaining 688 individuals. When also setting the score threshold for cancer samples to > 0.2, 98.9% of the 647 remaining samples could be matched correctly (Figure 4). This comparison suggested a high accuracy rate in population assignment for cancer samples and an increase in the level of accuracy with a lower admixture background of the individual.

**Fig. 4.**
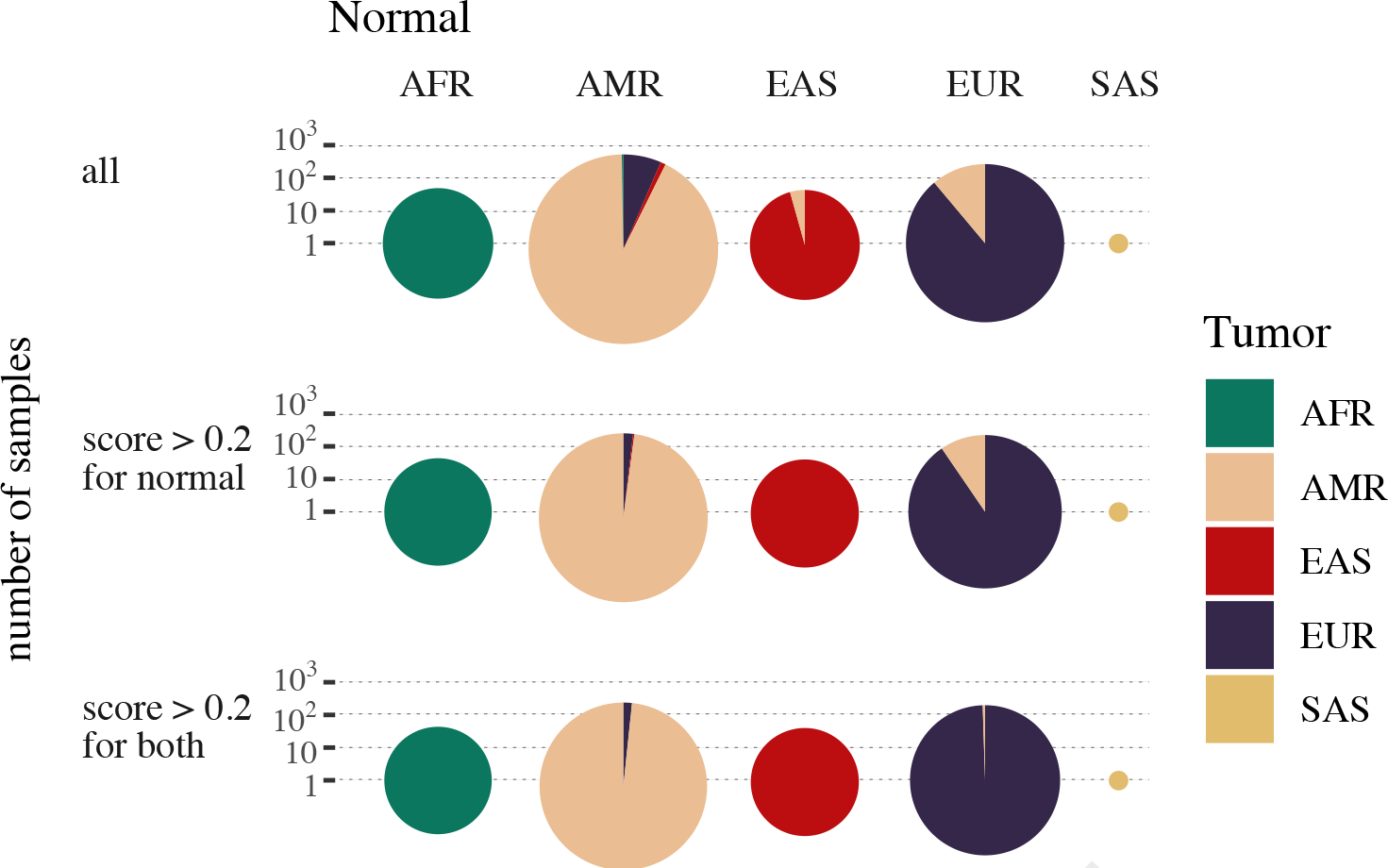
Accuracy of assignment with paired tumor and normal samples from GEO. 1145 individuals from GEO with paired tumor and normal samples were examined. First row of pie charts (“all”) indicates results of a total of 1145 individuals. Second row of pie charts (“score > 0.2 for normal)” indicates results for 688 individuals whose normal samples have scores higher than 0.2. Last row of pie charts (“score > 0.2 for both “) indicates results for 647 individuals, of whom both normal and cancer samples have scores higher than 0.2. The radius of the pie chart indicates the total number of samples. The columns are the prediction groups for the normal samples and the color codes indicate the proportion of each group predicted from their respective cancer samples.

#### TCGA data

We used the genomic array profiling data from TCGA project, all of which originate from the array platform Genome Wide SNP 6. We extracted all 18380 samples (of 9190 individuals), where a normal tissue control for the respective tumor sample was available. 8924 out of 9190 (97.1%) individuals had matched tumor/normal categories. After setting a threshold of normal samples with score > 0.2, 98.4% accuracy (8413 out of 8549 individuals) can be achieved. When also setting the score threshold for cancer samples to > 0.2, 8195 of the 8216 (99.7%) remaining samples could be matched correctly (Figure 5). The score and error relation was depicted in the Supplementary Table S5. For population assignment into 26 groups, 8522 out of 9190 (92.7%) pairs had matching categories (Supplementary Figure S4). With the same score thresholding as described above, the accuracy increased to 97% (7374 out of 7602) and 99.1% (6948 out of 7010) respectively.

**Fig. 5.**
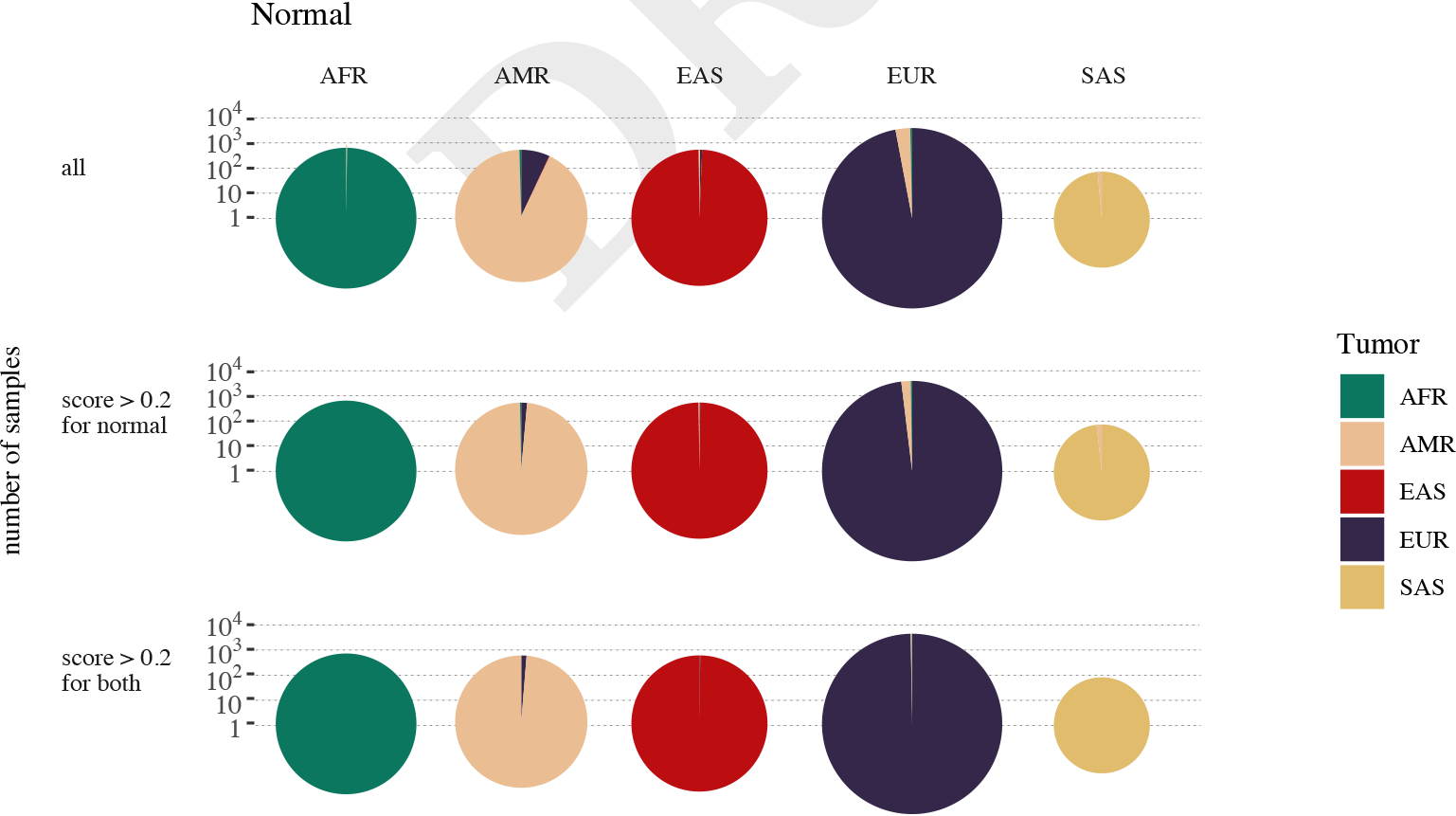
Accuracy of assignment with paired tumor and normal samples from TCGA project. 9190 individuals with paired tumor and normal samples from TCGA project were used for this benchmarking. First row of pie charts (“all”) indicates results of a total of 9190 individuals. Second row of pie charts (“score > 0.2 for normal)” indicates results for 8549 individuals whose normal samples have scores higher than 0.2. Last row of pie charts (“score > 0.2 for both “) indicates results for 8216 individuals, of whom both normal and cancer samples have scores higher than 0.2. The radius of the pie chart indicates the total number of samples. The columns are the prediction groups for the normal samples and the color codes indicate the proportion of each group predicted from their respective cancer samples.

#### Modification of SNP status during carcinogenesis

We further scrutinized the source of noise and information loss during carcinogenesis. Using the same dataset, we performed a SNP-by-SNP comparison between the paired normal and cancer samples and summarized the SNPs which changed from heterozygous to homozygous (Het2Homo), from homozygous to heterozygous (Homo2Het) or stayed identical (Supplementary Figure S5). The Het2Homo (6.3% on average) transition occurred in case of allele loss or copy-neutral loss of heterozygocity and constituted the larger part to noise in cancer samples. The Homo2Het (5.6% on average) transition could come from loss of both alleles causing BAF to appear at 0.5 or less frequently when somatic point mutations coincide with germline polymorphism sites. In this section, We benchmarked the cancer/normal assignment consistency with presence of these two types of SNP status modifications. We also indirectly addressed the issue of tumor sample purity here, as mixed cancer/normal samples would be assigned to same categories as either of the pure cell populations.

### Comparison with self-reported ethnicity metadata

To assess the overall accuracy of self-reported population information, we compared the meta-data from GEO and TCGA with our benchmarked results.

#### GEO data

We retrieved a total of 1724 samples with intelligible self-reported metadata from GEO. We extracted the population-implying keywords, which formed nine groups: “african”(92), “african-american”(59), “black”(6), “caucasian/european”(1472), “white”(40), “asian”(23), “chinese”(12), “hispanic”(12) and “native american”(2) (Figure 6). Specifically, “african” and “african-american” samples were mostly assigned to “AFR” group (91.3% and 93.6% respectively). The 1472 “caucasian/european”-labeled samples were assigned to “EUR” (90.2%) with small fraction assigned to “AFR”(4.6%) or “AMR”(5.0%). 11 out of 12 “hispanic” samples were assigned to “AMR” with one as “EUR”. All 40“white” samples were assigned to “AMR”. These 40 samples derived from the same study, so the patients were likely from a similar ethnicity background and reported as “white” by the study. The “asian” samples were mostly assigned to “EAS” group (19/23). All 12 “chinese” samples were correctly assigned to “EAS”. Two ‘native-american’ samples were both assigned to “EUR”. Counting all samples, a metadata matching rate of 88.3% was achieved. This indicated the existing heterogeneity in ethnicity description across studies and a certain degree of inaccuracy in self-reported ethnicity information.

**Fig. 6.**
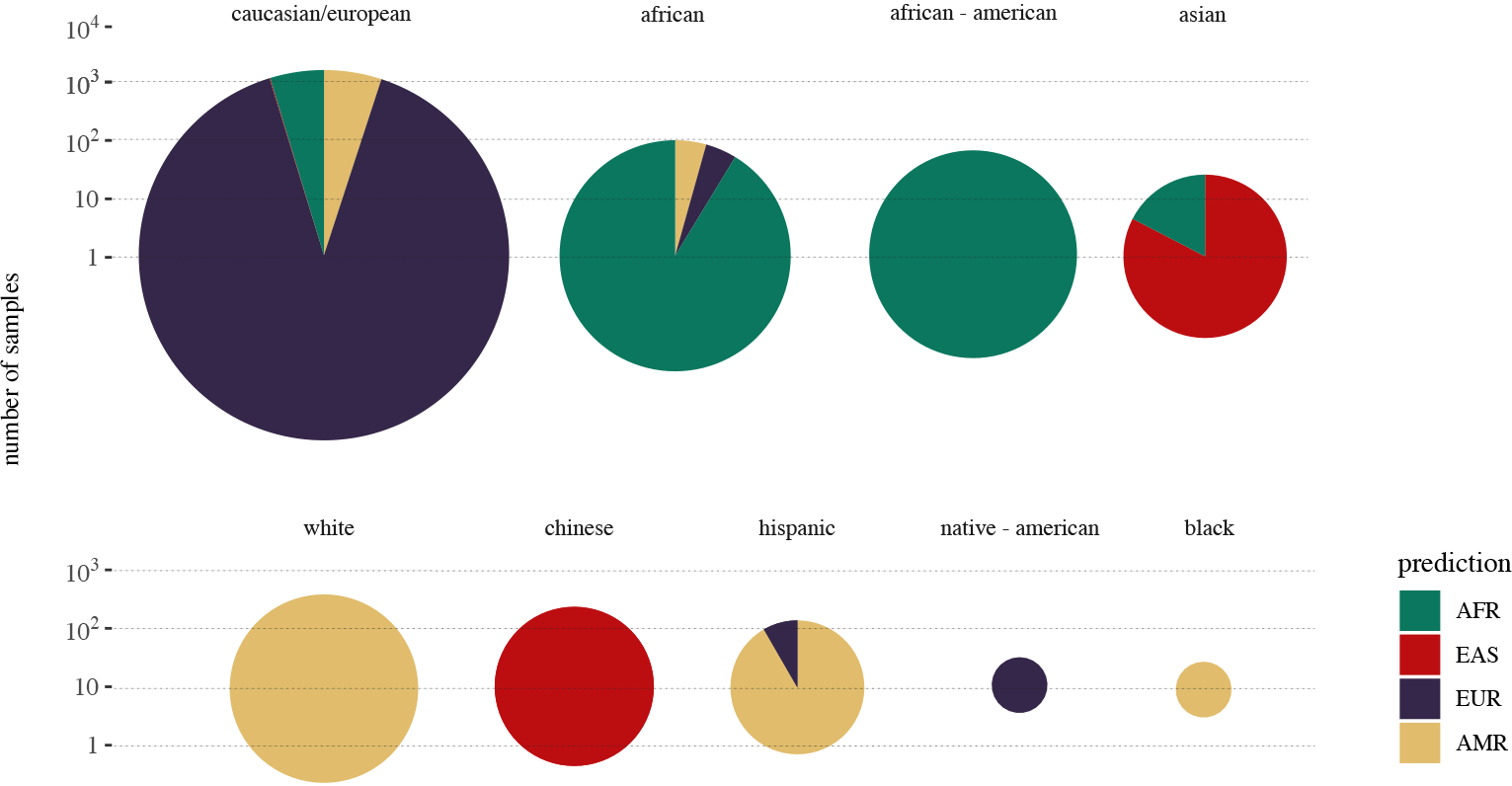
Accuracy of assignment with self-report metadata from GEO. The composition of assigned super population groups of 1724 samples from GEO with extracted population/ethnicity metadata in nine groups: “african”(92), “african-american”(59), “black”(6), “caucasian/european”(1472), “white”(40), “asian”(23), “chinese”(12), “hispanic”(12) and “native american”(2).

#### TCGA data

From the test of paired cancer/normal samples in TCGA, we also compared our results of 9190 normal samples with the “race” attribute provided by the metadata. There, six categories were distinguished: “American Indian or Alaska native”(23), “Asian”(612), “Black or African American”(793), “Native Hawaiian or other Pacific Islander”(12), “White”(6600) or “Not Reported”(1150). The fraction of assignment results in each of these categories were shown in the pie charts in Figure 7). Most samples reported as “white” were assigned to “EUR”(91.4%) with 7.7% “AMR” Most samples reported as “Black or African American” were assigned to “AFR” (95.6%) with “AMR” (3.0%) and EUR (1.0%); The “Asian”-labeled samples were composed of 90.5% “EAS” and 8.2% “SAS”. 14 out of the 23 “American Indian or Alaska Natives”-labeled samples were assigned to “AMR” and 5 to “EUR”. In the 12 “Native Hawaiian or other Pacific Islanders” samples, 10 were assigned to “EAS” and 2 to “AMR”. The samples were also assigned to one of the 26 population groups (Supplementary Figure S6), but there was no more meta-data for a comparison on this level. If we expect the following three“race” groups to match the assignment: “White” to “EUR”, “Black or African American” to “AFR” and “Asian” to “EAS” and “SAS”, then a total matching rate of 92.4% was achieved with 95.6% in African group, 98.6% in Asian group and 91.4% in European group. Despite the high matching percentage on the level of super population group in the self-report groups with large sample numbers (“White”, “Black or African American” and “Asian”), one may still argue that with the prediction outcome from our tool, the “race” information defined in the project could be well complemented and adapted for a quantitative measure for genetic background assessment.

**Fig. 7.**
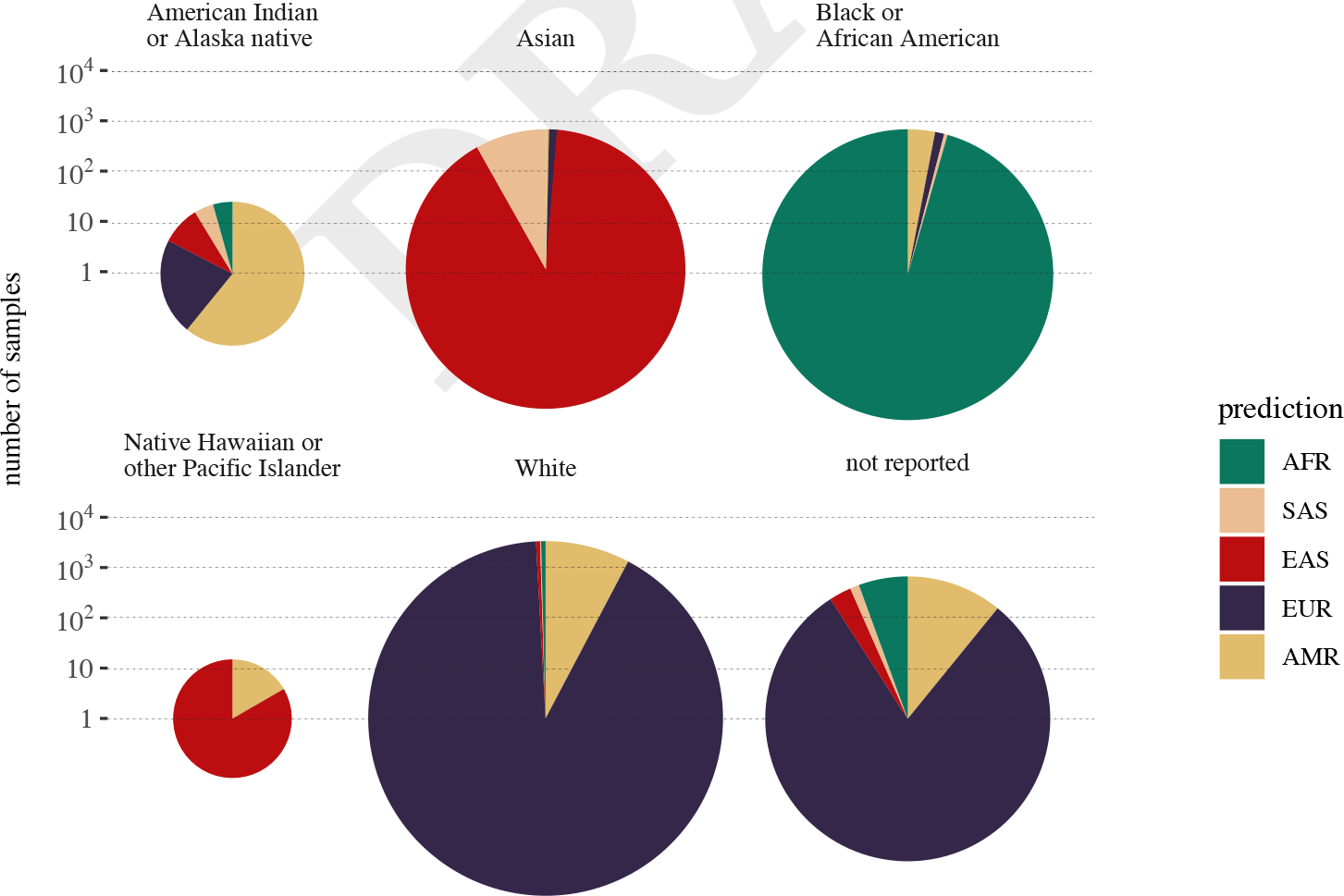
Accuracy of assignment with self-report metadata from TCGA project. The composition of assigned super population groups of 9190 samples in six “race” categories from TCGA meta-data: “American Indian or Alaska native”(23), “Asian”(612), “Black or African American”(793), “Native Hawaiian or other Pacific Islander”(12), “White”(6600) or “Not Reported”(1150).

## Discussion

The presented method resolves two issues that have been missing in the earlier approaches to estimate population structure from genotyping data: 1. it allows the integration of platforms with heterogeneous, non-overlapping SNP positions for cross-platform meta-analysis; 2. it confirms the possibility of deriving the population origin using data from aberrant cancer genomes in addition to those of unaltered genomes.

In our study, we establish the robustness of deriving population despite platform heterogeneity by cross-platform benchmarking. We show that for all the genotyping platforms used in the study, our pipeline achieves similarly high prediction accuracy, even for the lowest resolution platform with around 10 000 SNPs. This helps to integrate genotyping data generated with different platforms for meta-analysis, where the heterogeneity in resolution does not pose great impact on the parameters of interest (e.g. cancer copy number variation studies).

We demonstrate the feasibility of incorporating aberrant cancer genomes in a step-wise manner. First, we benchmark the method using samples with unaltered genomes (non-cancer) which are different from the reference dataset, and show a high matching accuracy. Then, we employ paired normal/cancer samples and benchmark the assignment consistency between them. With two independent data resources (GEO and TCGA), we show that the population group estimated from unaltered genome of the same individual matches that from the aberrant cancer genome with a high accuracy rate.

From the paired sample comparison, we do observe a certain degree of mismatches. These often occur in samples with a highly admixed background (indicated by lower score), rather than related to whether they are normal or cancer samples as discussed above. Furthermore, the mismatch occurs mostly between the AMR and EUR labels, which is partly expected as the 1000 Genomes Project describes “AMR” group as “Admixed American”. Clearly defining indigenous American population as reference is impractical due to the admixture events in the last hundreds of years in South America. Such admixture events inevitably pose a challenge when establishing stable reference groups for assessing the impact of population background on secondary phenotypes like mutational patterns in cancer. Finally, as many human populations are still limitedly represented or missing and major contribu-tions to human genetic variations are emerging (33), genomic population models will be amenable for re-assortment.

## Conclusions

We have developed a bioinformatics tool to assign population group based on SNP genotyping array data. We demonstrate its feasibility and accuracy on cancer samples, where somatic mutations can obfuscate part of the ancestry related SNP signal. This work can facilitate the re-analysis and meta-analysis of available cancer data by grouping samples with similar genetic background to study the potential genetic predisposition to cancer. In addition, our method provides the basis for subsequent haplotype phasing and refinement of genomic landscape for emerging somatic variation. With this tool, researchers will be able to integrate cancer genome profiling data from multiple resources to better assess the contribution of population background in population-specific mutation patterns occurring in cancer.

## Abbreviations

ACB: African Caribbeans in Barbados
AFR: African
ALDH2: Aldehyde Dehydrogenase 2 Family (Mitochondrial)
AMR: Admixed American
ASW: Americans of African Ancestry in SW USA
BAF: B allele frequency
BEB: Bengali from Bangladesh
CDX: Chinese Dai in Xishuangbanna, China
CEU: Utah Residents (CEPH) with Northern and Western European Ancestry
CHB: Han Chinese in Beijing, China
CHS: Southern Han Chinese
CLM: Colombians from Medellin, Colombia
EAS: East Asian
ESN: Esan in Nigeria
EUR: European
FIN: Finnish in Finland
Fst: Fixation Index
GBR: British in England and Scotland
GEO: Gene Expression Omnibus
GIH: Gujarati Indian from Houston, Texas
GWD: Gambian in Western Divisions in the Gambia
IBS: Iberian Population in Spain
ITU: Indian Telugu from the UK
JPT: Japanese in Tokyo, Japan
KHV: Kinh in Ho Chi Minh City, Vietnam
LD: linkage disequilibrium
LWK: Luhya in Webuye, Kenya
MAF: Minor allele frequency
MSL: Mende in Sierra Leone
MXL: Mexican Ancestry from Los Angeles USA
PCA: Principle Component Analysis
PEL: Peruvians from Lima, Peru
PJL: Punjabi from Lahore, Pakistan
PUR: Puerto Ricans from Puerto Rico
SAS: South Asian
SNP: Single nucleotide polymorphism
STU: Sri Lankan Tamil from the UK
TCGA: the Cancer Genome Atlas
TSI: Toscani in Italia
YRI: Yoruba in Ibadan, Nigeria

## Author’s contributions

QH developed the tool and performed benchmarking. MB conceived the project. Both authors contributed to data as-sembly and writing and editing of the manuscript.

## ACKNOWLEDGEMENTS

We thank Paula Carrio Cordo for the metadata curation and Bo Gao for technical support and helpful discussions. We thank the Zurich Bioinformatics group and ELIXIR Population Genomics for valuable comments.

## Supplementary Figures

**Fig. S1. The fraction or contribution of theoretical ancestors (k=9) in reference individuals from 1000 Genomes Project with regard to nine SNP array platforms.** The x-axis are individual samples, grouped by their respective population. Groups belonging to the same continent/superpopulation are placed neighboring to each other: AFR (1-7), SAS (8-12), EAS (13-17), EUR (18-22), AMR (23-26).

**Fig. S2. Percentage of accurate prediction with multinomial linear regression (MLR) model and random forest (RF) model into 26 population groups defined in 1000 Genomes Project.**

**Fig. S3. Percentage of accurate prediction with multinomial linear regression (MLR) model and random forest (RF) model into 10 population groups defined by the classification ambiguity between 26 groups. The frequently mixed groups are merged into one, leaving 10 out of 26 groups.** See Supplementary Table S3 for the merged groups.

**Fig. S4. Agreement of assigned population groups between paired tumor and normal samples in TCGA project on the level of 26 population groups.**

**Fig. S5. Percentage of SNPs in the genotyping arrays that stay identical or changed status (Homo2Het: homozygous to heterozygous or Het2Homo: heterozygous to homozygous) between paired cancer and normal samples.**

**Fig. S6. The composition of assigned population groups in six “race” categories from TCGA meta-data on the level of 26 population groups.**

## Supplementary Tables

**Table S1**. 12 IDs were removed from the reference set for the final prediction model due to high admixture background.

**Table S2**. Computation time per cross validation run with multinomial linear regression (MLR) model and random forest (RF) model with respect to the number of prediction output groups.

**Table S3**. 10 groups can be distinguished from each other, based on the platform-derived SNPs.

**Table S4**. Assignment concordance between paired samples in relation to assignment score range for 1145 paired cancer and normal samples from GEO.

**Table S5**. Assignment concordance between paired samples in relation to assignment score range for 9190 paired cancer and normal samples from TCGA.

## Bibliography

1. D Hanahan and RA Weinberg. Hallmarks of cancer: the next generation. Cell, 144(5): 646–674, 2011.

2. Ludmil B Alexandrov, Serena Nik-Zainal, David C Wedge, Samuel AJR Aparicio, Sam Behjati, Andrew V Biankin, Graham R Bignell, Niccolo Bolli, Ake Borg, Anne-Lise Børresen-Dale, et al. Signatures of mutational processes in human cancer. Nature, 500(7463):415, 2013.

3. HT Lynch and A de la Chapelle. Hereditary colorectal cancer. N Engl J Med, 348(10): 919–932, 2003.

4. J Zhang, KE Nichols, and JR Downing. Germline mutations in predisposition genes in pediatric cancer. N Engl J Med, 374(14):1391, 2016.

5. D Max Parkin, Paola Pisani, and J Ferlay. Global cancer statistics. CA: a cancer journal for clinicians, 49(1):33–64, 1999.

6. G Danaei, S Vander Hoorn, AD Lopez, CJ Murray, M Ezzati, and Risk Assessment collaborating group (Cancers Comparative. Causes of cancer in the world: comparative risk assessment of nine behavioural and environmental risk factors. Lancet, 366(9499):1784–1793, 2005.

7. RL Siegel, KD Miller, and A Jemal. Cancer statistics, 2017. CA Cancer J Clin, 67(1):7–30, 2017.

8. Rebecca L Siegel, Kimberly D Miller, and Ahmedin Jemal. Cancer statistics, 2018. CA: a cancer journal for clinicians, 68(1):7–30, 2018.

9. Laufey T Amundadottir, Patrick Sulem, Julius Gudmundsson, Agnar Helgason, Adam Baker, Bjarni A Agnarsson, Asgeir Sigurdsson, Kristrun R Benediktsdottir, Jean-Baptiste Cazier, Jesus Sainz, et al. A common variant associated with prostate cancer in european and african populations. Nature genetics, 38(6): 652, 2006.

10. Simon N Stacey, Andrei Manolescu, Patrick Sulem, Thorunn Rafnar, Julius Gudmundsson, Sigurjon A Gudjonsson, Gisli Masson, Margret Jakobsdottir, Steinunn Thorlacius, Agnar Helgason, et al. Common variants on chromosomes 2q35 and 16q12 confer susceptibility to estrogen receptor-positive breast cancer. Nature genetics, 39(7): 865, 2007.

11. Albert Tenesa, Susan M Farrington, James GD Prendergast, Mary E Porteous, Marion Walker, Naila Haq, Rebecca A Barnetson, Evropi Theodoratou, Roseanne Cetnarskyj, Nicola Cartwright, et al. Genome-wide association scan identifies a colorectal cancer susceptibility locus on 11q23 and replicates risk loci at 8q24 and 18q21. Nature genetics, 40(5):631, 2008.

12. Chen Wu, Zhibin Hu, Dianke Yu, Liming Huang, Guangfu Jin, Jie Liang, Huan Guo, Wen Tan, Mingfeng Zhang, Ji Qian, et al. Genetic variants on chromosome 15q25 associated with lung cancer risk in chinese populations. Cancer research, 69(12):5065–5072, 2009.

13. Steven A. Frank. Genetic predisposition to cancer - insights from population genetics. Nature Reviews Genetics, 5:764 EP –, Oct 2004. Review Article.

14. Yoshio Miki, Jeff Swensen, Donna Shattuck-Eidens, P Andrew Futreal, Keith Harshman, Sean Tavtigian, Qingyun Liu, Charles Cochran, L Michelle Bennett, Wei Ding, et al. A strong candidate for the breast and ovarian cancer susceptibility gene brca1. Science, 266(5182):66–71, 1994.

15. William D. Foulkes, Bartha Maria Knoppers, and Clare Turnbull. Population genetic testing for cancer susceptibility: founder mutations to genomes. Nature Reviews Clinical Oncology, 13:41 EP–, Oct 2015. Review Article.

16. Hui Li, Svetlana Borinskaya, Kimio Yoshimura, Nina Kal’ina, Andrey Marusin, Vadim A Stepanov, Zhendong Qin, Shagufta Khaliq, Mi-Young Lee, Yajun Yang, et al. Refined geographic distribution of the oriental aldh2* 504lys (nee 487lys) variant. Annals of human genetics, 73(3):335–345, 2009.

17. Philip J Brooks, Mary-Anne Enoch, David Goldman, Ting-Kai Li, and Akira Yokoyama. The alcohol flushing response: an unrecognized risk factor for esophageal cancer from alcohol consumption. PLoS medicine, 6(3):e1000050, 2009.

18. Tanya Keenan, Beverly Moy, Edmund A. Mroz, Kenneth Ross, Andrzej Niemierko, James W. Rocco, Steven Isakoff, Leif W. Ellisen, and Aditya Bardia. Comparison of the genomic landscape between primary breast cancer in african american versus white women and the association of racial differences with tumor recurrence. Journal of Clinical Oncology, 33(31):3621–3627, 2015. doi: 10.1200/JCO.2015.62.2126. PMID: 26371147.

19. Jiaying Deng, Hu Chen, Daizhan Zhou, Junhua Zhang, Yun Chen, Qi Liu, Dashan Ai, Hanting Zhu, Li Chu, Wenjia Ren, Xiaofei Zhang, Yi Xia, Menghong Sun, Huiwen Zhang, Jun Li, Xinxin Peng, Liang Li, Leng Han, Hui Lin, Xiujun Cai, Jiaqing Xiang, Shufeng Chen, Yihua Sun, Yawei Zhang, Jie Zhang, Haiquan Chen, Shijian Zhang, Yi Zhao, Yun Liu, Han Liang, and Kuaile Zhao. Comparative genomic analysis of esophageal squamous cell carcinoma between asian and caucasian patient populations. Nature Communications, 8(1):1533, 2017. ISSN 2041-1723. doi:10.1038/s41467-017-01730-x.

20. Wensheng Zhang, Andrea Edwards, Erik K. Flemington, and Kun Zhang. Racial disparities in patient survival and tumor mutation burden, and the association between tumor mutation burden and cancer incidence rate. Scientific Reports, 7(1): 13639, 2017. ISSN 2045-2322. doi:10.1038/s41598-017-13091-y.

21. Paul D. P. Pharoah, Antonis Antoniou, Martin Bobrow, Ron L. Zimmern, Douglas F. Easton, and Bruce A. J. Ponder. Polygenic susceptibility to breast cancer and implications for prevention. Nature Genetics, 31:33 EP –, Mar 2002. Article.

22. Anders Albrechtsen, Thorfinn Sand Korneliussen, Ida Moltke, Thomas van Overseem Hansen, Finn Cilius Nielsen, and Rasmus Nielsen. Relatedness mapping and tracts of relatedness for genome-wide data in the presence of linkage disequilibrium. Genetic epidemiology, 33(3):266–274, 2009.

23. Rust Turakulov and Simon Easteal. Number of snps loci needed to detect population structure. Human heredity, 55(1):37–45, 2003.

24. A Auton, LD Brooks, RM Durbin, EP Garrison, HM Kang, JO Korbel, JL Marchini, S McCarthy, GA McVean, GR Abecasis, and 1000 Genomes Project Consortium. A global reference for human genetic variation. Nature, 526(7571):68–74, 2015.

25. DH Alexander, J Novembre, and K Lange. Fast model-based estimation of ancestry in unrelated individuals. Genome Res, 19(9):1655–1664, 2009.

26. Petr Danecek, Adam Auton, Goncalo Abecasis, Cornelis A Albers, Eric Banks, Mark A DePristo, Robert E Handsaker, Gerton Lunter, Gabor T Marth, Stephen T Sherry, et al. The variant call format and vcftools. Bioinformatics, 27(15):2156–2158, 2011.

27. Shaun Purcell, Benjamin Neale, Kathe Todd-Brown, Lori Thomas, Manuel AR Ferreira, David Bender, Julian Maller, Pamela Sklar, Paul IW De Bakker, Mark J Daly, et al. Plink: a tool set for whole-genome association and population-based linkage analyses. The American Journal of Human Genetics, 81(3):559–575, 2007.

28. Maria Ortiz-Estevez, Henrik Bengtsson, and Angel Rubio. Acne: a summarization method to estimate allele-specific copy numbers for affymetrix snp arrays. Bioinformatics, 26(15): 1827–1833, 2010.

29. H Cai, N Kumar, and M Baudis. arraymap: a reference resource for genomic copy number imbalances in human malignancies. PLoS One, 7(5):e36944, 2012.

30. R Edgar, M Domrachev, and AE Lash. Gene expression omnibus: Ncbi gene expression and hybridization array data repository. Nucleic Acids Res, 30(1):207–210, 2002.

31. Cancer Genome Atlas Research Network. Comprehensive genomic characterization defines human glioblastoma genes and core pathways. Nature, 455(7216):1061–1068, 2008.

32. International HapMap Consortium. The international hapmap project. Nature, 426(6968): 789–796, 2003.

33. Rachel M. Sherman, Juliet Forman, Valentin Antonescu, Daniela Puiu, Michelle Daya, Nicholas Rafaels, Meher Preethi Boorgula, Sameer Chavan, Candelaria Vergara, Victor E. Ortega, Albert M. Levin, Celeste Eng, Maria Yazdanbakhsh, James G. Wilson, Javier Marrugo, Leslie A. Lange, L. Keoki Williams, Harold Watson, Lorraine B. Ware, Christopher O. Olopade, Olufunmilayo Olopade, Ricardo R. Oliveira, Carole Ober, Dan L. Nicolae, Deborah A. Meyers, Alvaro Mayorga, Jennifer Knight-Madden, Tina Hartert, Nadia N. Hansel, Marilyn G. Foreman, Jean G. Ford, Mezbah U. Faruque, Georgia M. Dunston, Luis Caraballo, Esteban G. Burchard, Eugene R. Bleecker, Maria I. Araujo, Edwin F. Herrera-Paz, Monica Campbell, Cassandra Foster, Margaret A. Taub, Terri H. Beaty, Ingo Ruczinski, Rasika A. Mathias, Kathleen C. Barnes, and Steven L. Salzberg. Assembly of a pangenome from deep sequencing of 910 humans of african descent. Nature Genetics, 2018. ISSN 1546-1718. doi:10.1038/s41588-018-0273-y.

