## supplementary figures 1-6 for "Enabling population assignment from cancer genomes with SNP2pop"

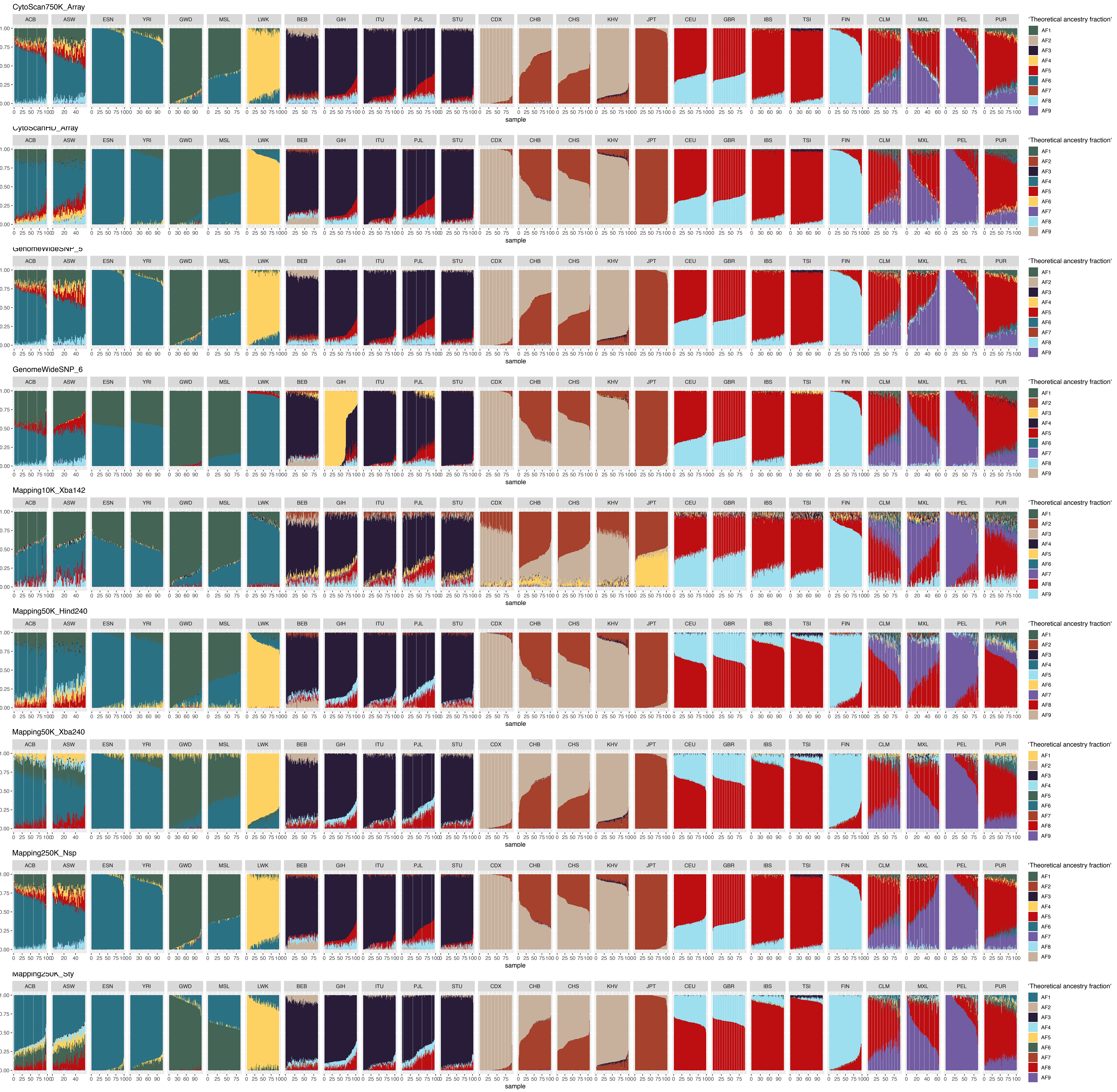


Supplementary Figure 1

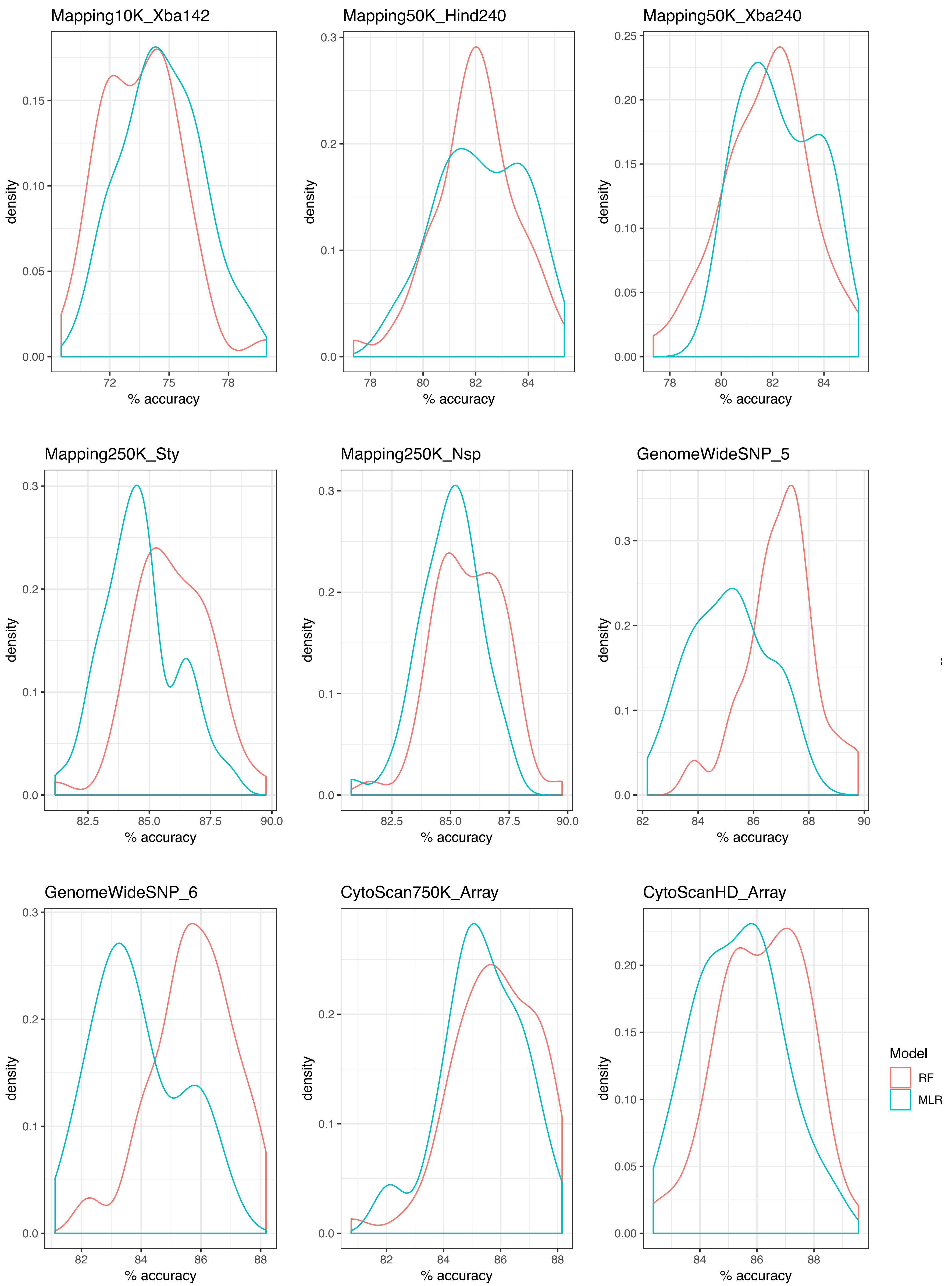


Supplementary Figure 2

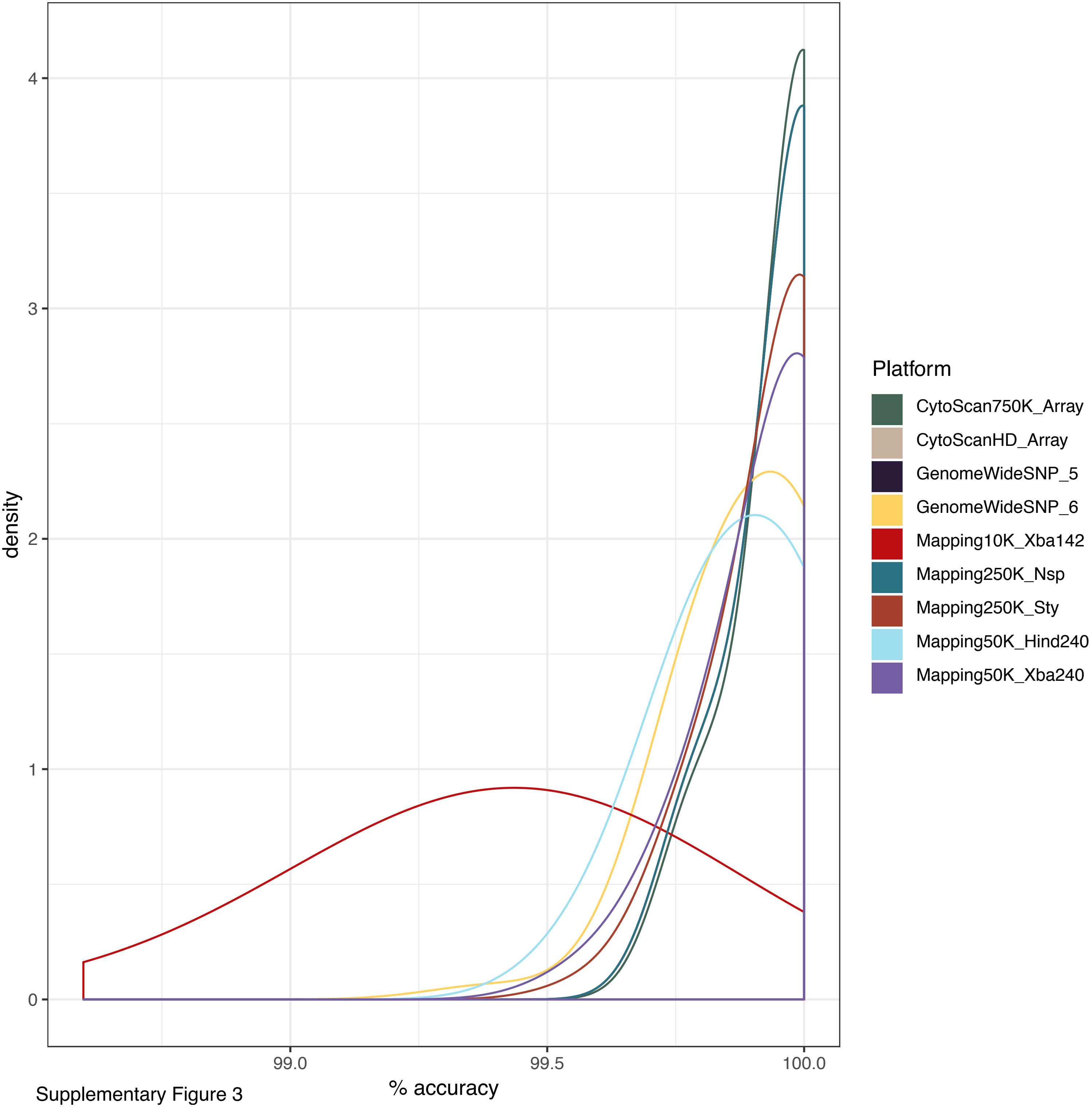


Supplementary Figure 3

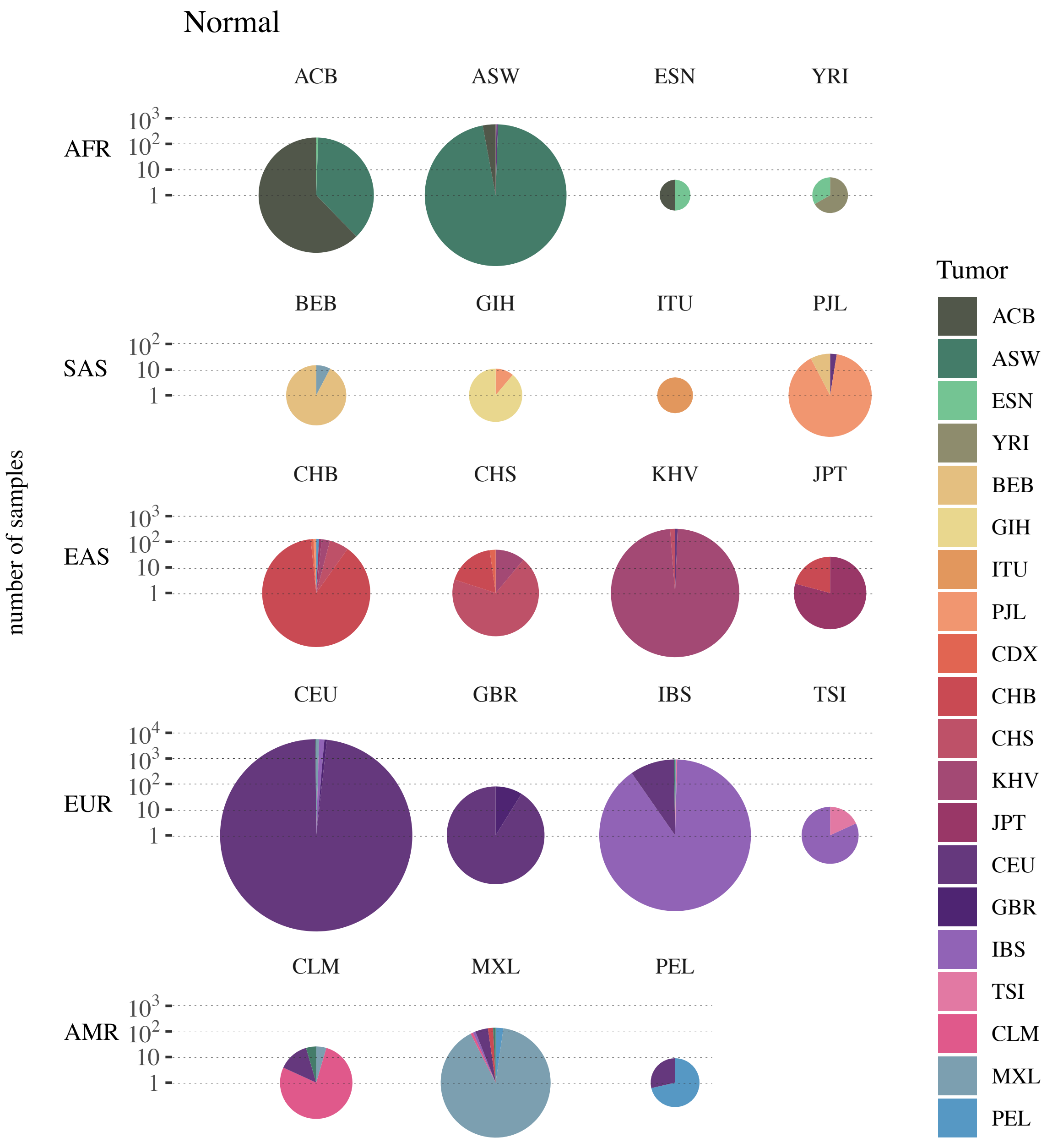


Supplementary Figure 4

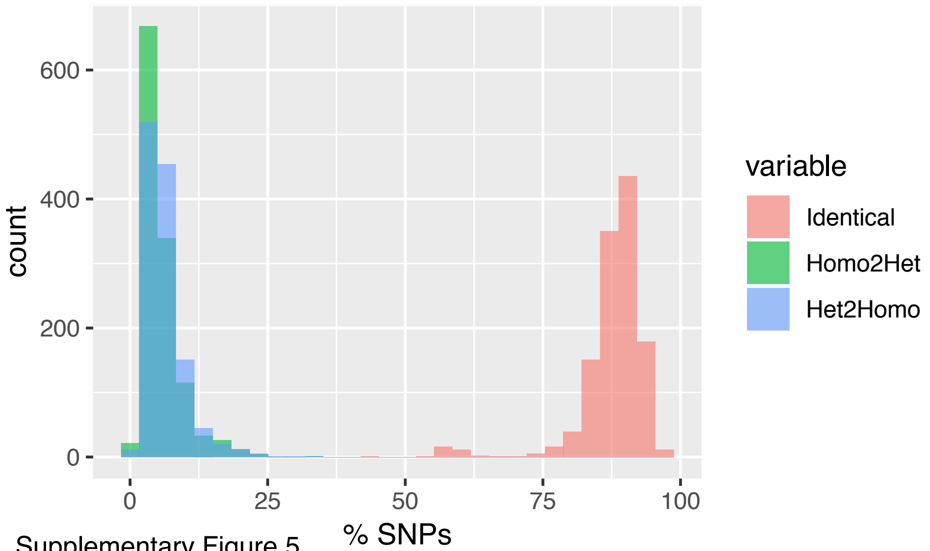


Supplementary Figure 5

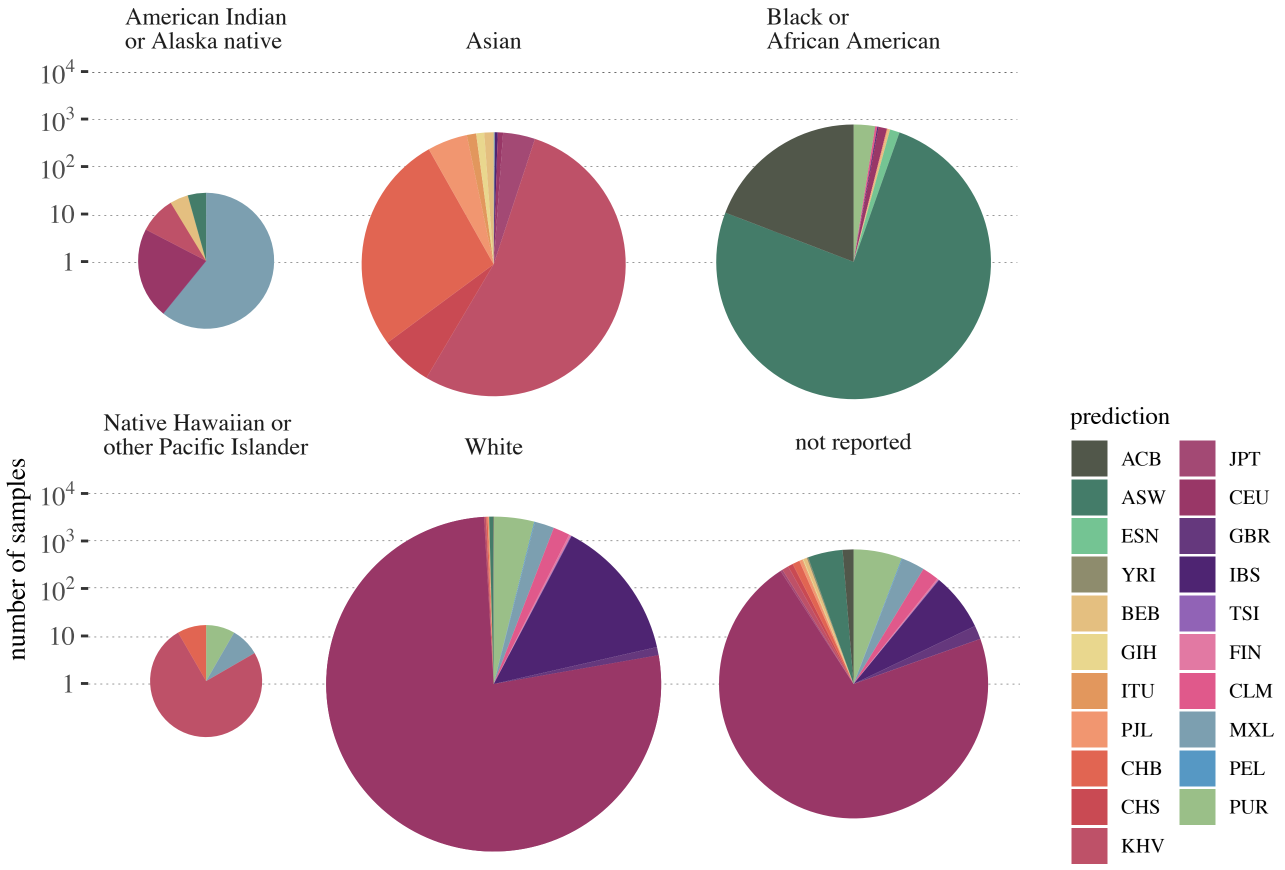


Supplementary Figure 6
